# The use of barcoded *Asaia* bacteria in mosquito *in vivo* screens for identification of systemic insecticides and inhibitors of malaria transmission

**DOI:** 10.1101/2021.09.29.462277

**Authors:** Angelika Sturm, Martijn W. Vos, Rob Henderson, Maarten Eldering, Karin M.J. Koolen, Avinash Sheshachalam, Guido Favia, Kirandeep Samby, Esperanza Herreros, Koen J. Dechering

## Abstract

This work addresses the need for new chemical matter in product development for control of pest insects and vector-borne diseases. We present a barcoding strategy that enables phenotypic screens of blood-feeding insects against small molecules in microtiter plate-based arrays and apply this to discovery of novel systemic insecticides and compounds that block malaria parasite development in the mosquito vector. Encoding of the bloodmeals was achieved through recombinant DNA-tagged *Asaia* bacteria that successfully colonized *Aedes* and *Anopheles* mosquitoes. An arrayed screen of a collection of pesticides showed that chemical classes of avermectins, phenylpyrazoles and neonicotinoids were enriched for compounds with systemic adulticide activity against *Anopheles*. Using a luminescent *Plasmodium falciparum* reporter strain, barcoded screens identified 48 drug-like transmission blocking compounds from a 400-compound antimicrobial library. The approach significantly increases the throughput in phenotypic screening campaigns using adult insects, and identifies novel candidate small molecules for disease control.

## Introduction

Parasites and viruses that are carried by mosquitoes cause diseases such as malaria, dengue or yellow fever. Malaria resulted in 229 million cases causing 409.000 deaths in 2019. (WHO World malaria report 2020). The use of insecticides has had large impact on control of malaria (*1*). Since World War II, the range of chemical scaffolds with insecticide activity has slowly expanded resulting in 55 chemically distinct classes of marketed insecticides available in 2019(*2*). Concurrently, resistance to these molecules has developed at a similar rate as a result of wide-spread use in crop protection, community and household spraying and impregnation of bed-nets (*3*). As a more targeted approach, the use of oral insecticides in drug-based vector control is considered (*4*). The endectocide ivermectin is used as an oral helminticide but also shows systemic adulticide activity against *Anopheles* mosquitoes (*5*). It has shown promise as a drug that, following repeat mass drug administration to a human population at risk, reduces malaria burden by directly blocking onward pathogen transmission through reduction of the life span of blood-feeding mosquitoes (*6*). Ivermectin is relatively rapidly eliminated from the blood circulation in humans, whereas modelling suggest that the duration of the mosquitocidal activity strongly drives impact of drug-based vector control (*7*). Therefore, long-acting drug substances and formulations are being pursued (*8, 9*).

As an alternative to use of insecticides for control of vector-borne disease, strategies aimed at biological control of the pathogen stages that underlie spread of the disease are emerging. These approaches have the advantage of a low risk on development of resistance. Arboviruses like zika and dengue and protozoa such as *Leishmania, Plasmodium* and *Trypanosoma* face a population bottleneck in the insect vector(*10, 11*) with a low number of replication cycles and, hence, a low rate of accumulation of resistance mutations. In the context of malaria elimination, drug interventions targeting the transmission stages of the malaria parasite are explored (*12*). Such compounds may kill or sterilize sexual stage parasites that infect mosquitoes (*13, 14*). Historically, antimalarial compounds have been selected on their ability to clear asexual blood stage parasitaemia that underlies clinical disease, and many of these compounds do not block transmission. More recently, compounds have emerged with a transmission-blocking component in their activity spectrum, although in many cases this activity is not as potent as their activity against asexual blood stages (*15, 16*). Therefore, there is a need for novel chemical starting points for development of malaria transmission-blocking drugs.

The requirements for drug candidates that block malaria transmission by killing the mosquito vector or by targeting the sexual stage parasites are outlined in target candidate profiles (TCP) 5 and 6, as put forward by the Medicines for Malaria Venture (*17*). These TCPs are stimulating and guiding global drug discovery efforts (*18*). In the absence of a large array of validated molecular targets, these efforts rely on phenotypic screens that have a relative low throughput and, hence, generate low numbers of chemically diverse starting points (*2, 19, 20*). In pesticide discovery, miniaturized assays in 96 wells assays containing larvae are used to predict systemic activity against adult insects (*21, 22*). Discovery of molecules that block transmission of malaria ultimately relies on laborious membrane-feeder experiments that use one container of mosquitoes for each test condition (*23*). An increase in throughput of these technologies would accelerate the development of novel malaria interventions. Here we present a technique that significantly improves the throughput of compound testing in phenotypic assays using adult mosquitoes. It allows screening of multiple molecules using barcoded bloodmeals in multi sample arrays. We used a genetically engineered prokaryotic symbiont, α-Proteobacteria of the genus *Asaia*, that stably associate with a number of sugar feeding insects(*24*). Upon ingestion with a glucose or a blood meal *Asaia* actively colonises the insect midgut within one or two days and spreads from there to most other organs(*25, 26*). We transformed *Asaia* strains with plasmids that carry individual short DNA barcodes. Following feeding of mosquitoes on arrays of bloodmeals with test compounds, these DNA-barcodes were recovered from the mosquito in order to deconvolute the feeding pattern and identify active compounds.. We used this technique to identify systemic insecticides and malaria transmission-blocking compounds from libraries of small molecules.

## Results

### Membrane-feeding mosquitoes with a barcoded bloodmeal

We envisaged to use recombinant insect midgut bacteria as tag bloodmeals in 96 well plates presented to hematophagous insects, allowing deconvolution of the feeding pattern. A previous study has shown the feasibility of feeding *Anopheles* mosquitoes on 96 well plates covered with Parafilm membrane (*27*). We developed a device to evenly stretch a Parafilm membrane in two directions (Fig. S1a). A hydraulic press was used to push the stretched membrane firmly down on a 96 well microtiter plate containing prewarmed bloodmeals (Fig. S1b). The plate was placed upside down on a cage of mosquitoes and warmed with a pre-heated aluminium heat block that was routed to exactly fit the base of the microtiter plate (Fig. S1c). Feeding efficiency depended on the temperature of the heat block. At 45 °C feeding performance was consistently above 90%, which was comparable or even better than a method using conventional glass feeders (Fig. 1a). Video analysis of feeding behaviour on a cage with ∼300 mosquitoes suggested sampling of all wells across the plate (Suppl. Video1).

**Figure 1.**
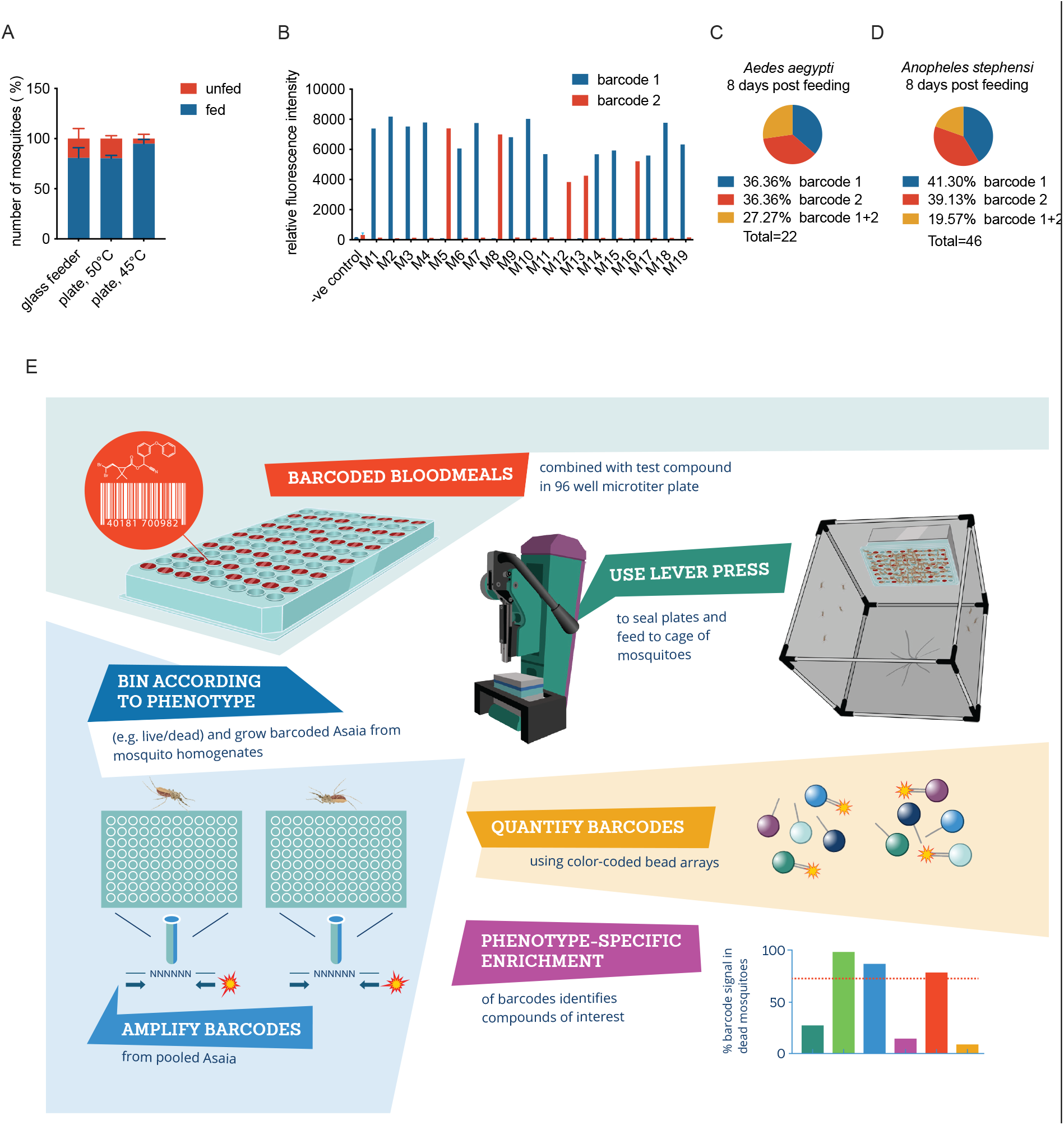
Tagging mosquitoes with a molecular barcode through feeding on microtiter plates. A) Comparison of feeding efficiency of *Anopheles stephensi* mosquitoes between glass feeders and microtiter plates. An aluminium block heated to 50 or 45 °C as indicated on the x-axis was placed on top of the plate. The figure shows the percentage of fed and unfed mosquitoes in the cage. B) Barcode signals in individual *Aedes aegypti* mosquitoes fed on a grid containing two different barcodes. The mosquitoes were analysed 2 days after feeding and the figure shows relative fluorescence intensities for the two barcodes in individual mosquito samples. C) Prevalence of barcode positive *Ae. aegypti* mosquitoes 8 days after feeding on a grid of 2 distinct barcodes. D) Prevalence of barcode positive *An. stephensi* mosquitoes 8 days after feeding on a grid of 2 distinct barcodes. E) Outline of screening strategy for identification of systemic insecticides. Mosquitoes were fed on microtiter plates containing barcoded bloodmeals supplemented with test compounds. Two days after blood feeding, mosquitoes were split into pools of live and dead mosquitoes, and *Asaia* bacteria were grown from homogenates of individual mosquitoes in 96 well liquid cultures under kanamycin selection pressure. Barcodes were then amplified by PCR using a fluorescently labelled primer pair that binds a common sequence flanking the DNA-barcode sequence. Following amplification, barcodes were quantified by multi analyte profiling using DNA oligos coupled to color-coded microspheres(*28*), which resulted in a fluorescence signal for each barcode depending on the quantity of the barcode in the PCR amplification product. Barcodes enriched in the dead mosquitoes identified compounds with systemic adulticidal activity, whereas detection of barcode signals from the live mosquitoes were used to verify sampling of barcodes that were missing in the pool of dead mosquitoes.

For introduction of a molecular barcode into mosquitoes, we evaluated two potential bacterial carriers, the midgut symbionts *Pantoe agglomerans* and *Asaia SF2*.*1* (*26, 29*). *Pantoea* showed a strong effect on transmission of *Plasmodium falciparum* malaria parasites to *Anopheles stephensi* mosquitoes (Fig. S2) and all subsequent experiments used *Asaia* SF2.1. We generated a collection of 50 bacterial stocks each with a unique DNA tag (Tables S1-S3). Pilot experiments with *Aedes aegypti* mosquitoes fed on a grid with two differently barcoded bloodmeals each placed in three different wells in a 96 well plate showed that 2 days after feeding, 100 percent of the fed mosquitoes was successfully tagged with a single barcode (Fig. 1b). Out of these, 74% contained barcode 1 and 26% contained barcode 2. This uneven distribution of barcodes may relate to the relatively small sample size. Analyses of a different cohort of mosquitoes 8 days after feeding showed a more even distribution, with equal proportions (36%) of mosquitoes having a single barcode (Fig. 1c). At this timepoint, 27% showed a signal for both barcodes. Cross-feeding was not observed for the cohort analysed 2 days post-feeding. Alternatively, cross-contamination may occur later on in the experiment, possibly through contact with mosquito diuresis fluids, excrements or the cotton pad that was used for glucose feeding during the experiment. Pilot experiments with *An. stephensi* mosquitoes showed similar results, with roughly equal proportions of mosquitoes with a single barcode and 20% of mosquitoes with two barcodes (Fig. 1d). To prevent cross-contamination of barcodes in subsequent experiments, we limited exposure to glucose pads to two hours per day while changing pads daily. In addition, mosquitoes were transferred to new cages directly after feeding to reduce exposure to diuresis fluid on the cage floor.

### Phenotypic screen for systemic insecticide activity

Based on these initial pilot experiments, we devised a strategy for multiplex detection of barcode signals in order to enable larger phenotypic screens (Fig. 1e). We explored suitability of this screening principle for phenotypic screening for systemic insecticides using fipronil as a reference compound. *An. stephensi* mosquitoes were fed on a 96-well plate with 24 barcoded bloodmeals, half of them containing 10 μM fipronil and the other half 0.1% DMSO as a vehicle control (0.1% DMSO). 48 hours after feeding, we retrieved all blood-fed mosquitoes from the cage. Of these, 70 of were alive and 77 were dead. Analyses of barcode presence in individual mosquitoes showed that 100% of the mosquitoes were successfully tagged with a barcode. Of these, 124 (84.3%) showed a single barcode, 21 (14.3%) showed two barcodes, and two mosquitoes (1.4%) showed three barcodes (Fig. 2a).

**Figure 2.**
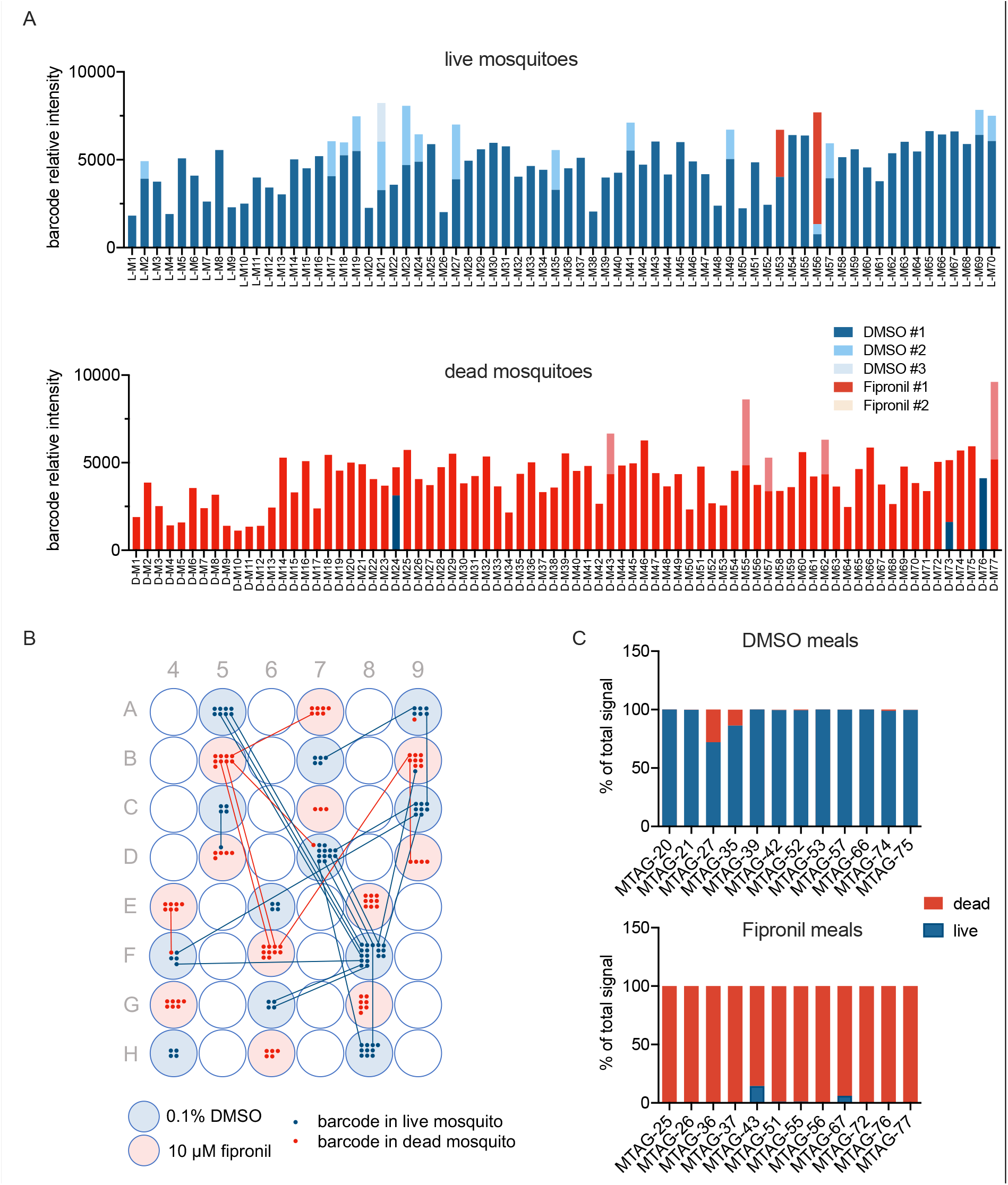
Proof of principle for systemic insecticide screen. *An. stephensi* mosquitoes were fed on a grid of 12 individually barcoded vehicle control (0.1% DMSO) and 12 individually barcoded insecticide (10 μM fipronil) bloodmeals. Two days after feeding, phenotype (live/dead) and barcode presence was determined for individual mosquitoes. A) Barcode signals in the live (upper panel) and dead (lower panel) cohort of mosquitoes. The figure shows the sum of barcode signals in individual mosquitoes. For mosquitoes showing multiple signals, barcode signals were grouped according to origin (DMSO or fipronil) and ranked on signal strength. DMSO #1, #2, #3 indicates the highest, second highest and third highest signal, respectively, originating from a DMSO well. There were no mosquitoes with more than three barcode signals above background. For the fipronil barcodes, we did not observe any mosquito with more than two barcodes originating from a fipronil-containing well. B) Deconvolution of feeding/cross-contamination pattern. Blue and red shading indicates wells that contained a DMSO or a fipronil bloodmeal, respectively. Dots indicate mosquitoes found positive for a barcode originating from that well, with blue dots indicating the mosquito was alive whereas red dots indicate dead mosquitoes. Lines indicate mosquitoes that were found positive for more than one barcode. C) Analyses of pooled samples. *Asaia* rescued from the mosquito midguts were binned according to the mosquito phenotype (live or dead) and barcodes were amplified from the pooled samples. The graphs show the proportion of barcode signal originating from live or dead mosquitoes for DMSO (upper panel) or fipronil barcoded bloodmeals. Barcodes indicated on the x-axis are listed in Table S3.

The barcodes associated with DMSO and fipronil segregated with a live and death phenotype, respectively. In the live cohort, two mosquitoes were positive for both DMSO and fipronil associated barcodes (Fig. 2a). This may be explained by intake of a sublethal quantity of fipronil, or cross-contamination of barcodes post feeding. In the dead cohort, all mosquitoes except for one were found positive with a fipronil-associated barcode. Deconvolution of the feeding pattern showed that wells were sampled on average by 7 mosquitoes, with a range of 3 to 17 mosquitoes (Fig. 2b). Cross-contamination of barcodes, either by uptake of multiple bloodmeals or post-feeding contact with barcode-containing material, was low across the plate. Since every well was sampled by multiple mosquitoes, the contaminating signal may only make up a small contribution to the summed signals from that well. For each barcode, we calculated the total signal from all mosquitoes positive a particular barcode in live or dead mosquitoes. The results show a clear compound and phenotype-dependent enrichment of barcodes (Fig. S3) and suggest that analyses of pooled samples may correctly annotate activity of a test compound in a barcoded bloodmeal. To test this experimentally, we created pools of *Asaia* bacteria rescued from live and dead mosquitoes, respectively, and analysed barcode intensities per pool in a single multiplex reaction. Barcodes in DMSO control meals were predominantly found in live mosquitoes, whereas barcodes in fipronil-containing bloodmeals were associated with the dead phenotype (Fig. 2c). The highest contaminated signal was observed with barcode 27 that was associated with a DMSO containing bloodmeal but showed 28% of the total signal originating from dead mosquitoes (Fig. 2c). This barcode was located in well A9, and was retrieved from a total of seven mosquitoes, of which 1 mosquito was dead at the time of sampling (Fig. 2b). The combined data highlight the feasibility of multiplex barcode detection in pools of mosquitoes binned according to the phenotype of interest.

### Screening pesticides against Anopheles stephensi mosquitoes

The above experiments demonstrated the feasibility of multiplex barcode detection in pools of mosquitoes binned according to the phenotype of interest. Using this strategy, we screened a collection of 83 chemically diverse pesticides to identify novel candidates for drug-based vector control approaches. Compounds were initially tested at 1 μM in duplicate with up to 48 samples per plate (Fig. S4). For each phenotype (live or dead mosquitoes), the *Asaia* cultures were pooled and DNA-barcodes from each pool were amplified and quantified. All compounds with ≥50% of the barcode signal in the dead mosquitoes were subsequently tested at 100 nM, whereas all inactive compounds (<50%) were tested at 10 μM concentration. From a total of 189 experimental conditions in 4 feeding experiments a total of 2727 mosquitoes was analysed. Of these, 952 were dead 48 hours after feeding. 4 wells were not sampled. One of these contained DMSO whereas other DMSO-containing wells were sampled normally. Compounds MMV03891 and MMV1577456 were not sampled in the initial run when tested at 1 μM but showed a barcode signal when tested at the same concentration in a repeat experiment, suggesting the initial lack of sampling was not due to interference, e.g. through a gustatory effect preventing bloodfeeding or antimicrobial action against the barcoded *Asaia*. Compound MMV1633827 was sampled when tested at 1 μM but not at 10 μM. The latter concentration was not repeated and we cannot exclude that this compound interfered at some point in the process. Fig. 3a shows the barcode signals for the negative (DMSO) and positive (deltamethrin and fipronil) control wells. The data indicate a clear treatment-dependent distribution of barcode signals over the two phenotypes, with an average of 100% of the signal in the dead mosquitoes for barcodes associated with either one of the insecticides, and 0% for barcodes associated with the DMSO control wells. The barcode distribution for all 189 experimental conditions showed a similar pattern, with a subpopulation around 0% and another around 100% associated with the death phenotype (Fig. 3b). Fig. 3c and Table S4 show the data for individual compounds. For four compounds we tested two different chemical batches, listed under separate MMV batch codes. Of these, methomyl (MM003972-04 & MM003972-05), nitempyram (MMV673126-3 & MMV673126-4) and amitraz (MMV002471-05 & MMV002471-06) showed consistent results between the two batches. For rotenone, batch MMV002519-09 did show activity at 10 μM whereas batch MMV002519-11 did not. Compounds from the class of avermectins appeared to be among the most active compounds with more than 90% of the barcode signal associated with the death phenotype at test concentrations of 100 nM and 1 μM (Fig. 3c, S5). Likewise, phenypyrazoles fipronil and vaniliprole and the isoxazoline fluralaner showed potent killing activity. Other phenylpyrazoles showed less potent activity, with more than 95% of the signal in the dead pool of mosquitoes when tested at 1 μM but not at 100 nM. The class of neonicotinoids was also enriched among the set of active compounds, with 70-100% of the barcode signal associated with the death phenotype when tested at either 1 or 10 μM. To validate the results from the barcoded screen we tested a number of compounds in traditional glass feeder membrane feeding experiments. These experiments confirmed systemic insecticide activity for all compounds tested (fipronil, deltamethrin, chlorfenapyr, abamectin, fluralaner, vaniliprole. spinetoram), with IC_50_ values ranging from 3 nM for abamectin to 3173 nM for chlorfenapyr (Fig. 3d).

**Figure 3.**
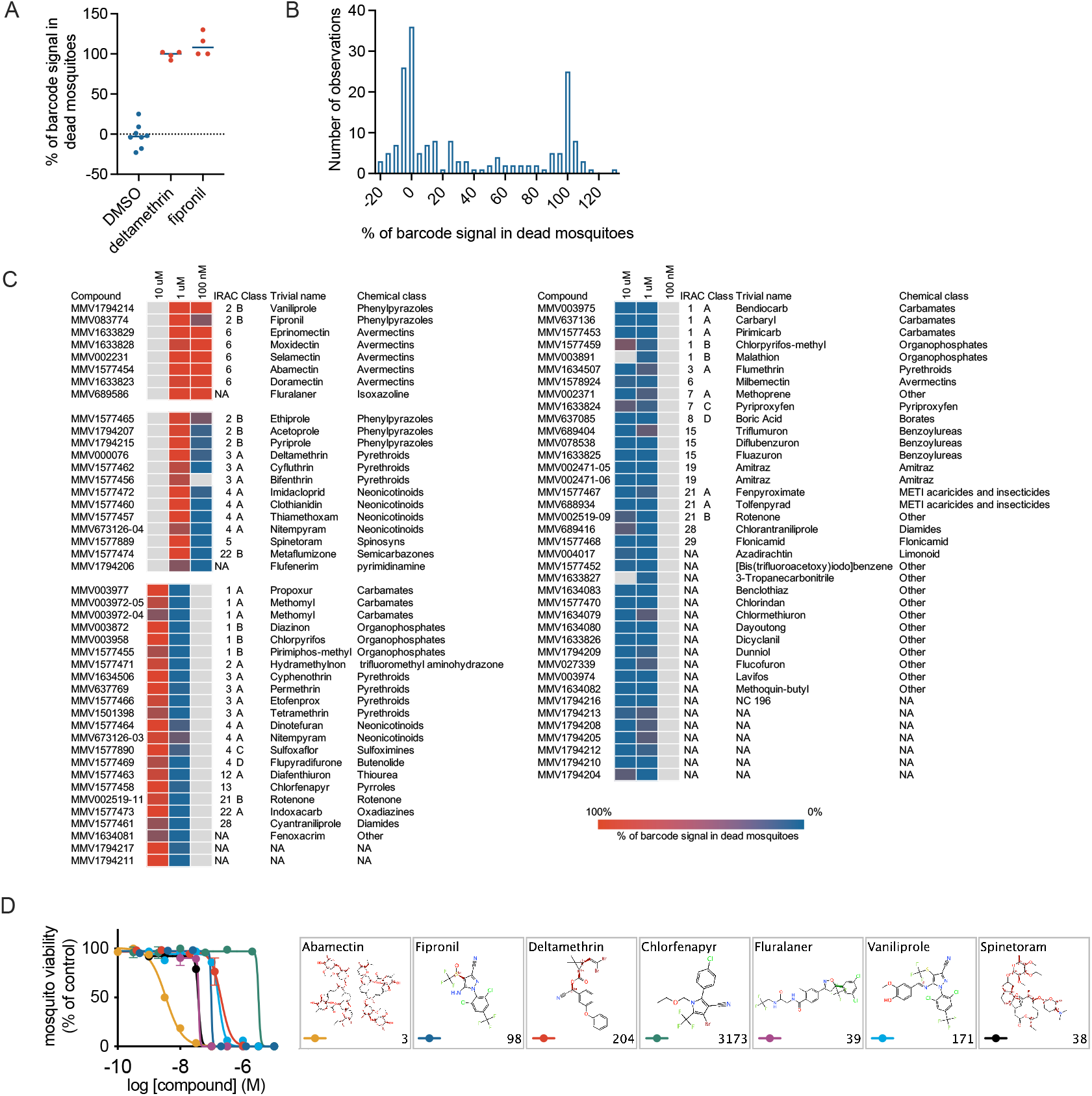
Screening of a collection of pesticides/insecticides. A) Assay controls. The figure shows the percentage of the total barcode signal that associated with the death phenotype for barcodes in control wells containing vehicle (0.1% DMSO), 10 μM deltamethrin or 10 μM fipronil. B) Distribution of barcode enrichment for all test conditions and barcodes. The figure shows a histogram of the proportion of the signal that was retrieved from dead mosquitoes relative to the total signal (dead plus live mosquitoes) for a particular barcode. C) Heatmap of barcode enrichment in dead mosquitoes for the compounds and test concentrations indicated. Compounds were initially tested at 1 μM. Inactive compounds were then tested at 10 μM whereas active compounds were tested at 100 nM. The color shading indicates the percentage of barcode enrichment in the dead mosquitoes. Grey colors indicate conditions that were not tested/sampled. D) Confirmation of systemic insecticide activity through traditional membrane feeding experiments using glass feeders. The compounds indicated in the legend were tested at multiple concentrations in duplicate. Error bars indicate standard deviations. IC_50_ estimates (in nM) from non-linear regression analysis are indicated in the lower right corners of the panels depicting the compound structures.

### Compound screen for transmission-blockade of Plasmodium falciparum

We developed a method for screening for inhibition of pathogen transmission, using the human malaria parasite as a model organism. In the procedure, outlined in Fig. S6, a transgenic *Plasmodium falciparum* reporter strain was used to infect *An. stephensi* mosquitoes by feeding on arrays of barcoded bloodmeals containing test compounds. This reporter produces a clear luminescence signal in infected mosquitoes eight days after feeding, when mature oocysts appear in the mosquito midgut (*30, 31*). At this time, mosquito infection status was assessed by luminescence measurement and *Asaia* bacteria were then rescued from individual mosquitoes and pooled into separate bins for infected and uninfected mosquitoes. Enrichment of barcode signals in the pool of uninfected mosquitoes identified wells containing a transmission-blocking test specimen. We screened the open access Pathogen Box, a collection of 400 chemically diverse and drug-like molecules selected for their potential action against a variety of pathogens underlying tropical infectious diseases (Fig. S7)(*32*). The total experiment involved 441 barcoded samples that were processed in 9 batches involving analyses of 4545 mosquitoes. Of these, 1794 showed a luminescence signal within 5 standard deviations of average background signal from unfed control mosquitoes and were considered uninfected (Fig. 4a). All barcodes were successfully detected in either uninfected, infected or both mosquito pools. For barcodes associated with atovaquone, on average 96% of the barcode signal was retrieved from the uninfected pool of mosquitoes (Fig. 4b). For the DMSO controls wells, the percentages of the barcode signals in the uninfected mosquitoes relative to the total barcode signals averaged at 18%. This is in line with (23)the experimental variation in mosquito infection success rates (*33*). Subsequently, we arbitrarily set the threshold for transmission-reducing activity at 80% of the barcode signal in the uninfected pool of mosquitoes, which separates the atovaquone from the DMSO vehicle controls with one exception (Fig. 4b). From the collection of 400 Pathogen Box compounds, 48 compounds met this criterion (Fig. 4c & Table S5). To verify this result, we selected 21 chemically diverse compounds for which barcodes were enriched in uninfected mosquitoes and tested these in individual membrane feeding experiments using regular glass feeders. Of these, 19 compounds reduced oocyst intensities by 80% or more in the glass feeder experiments, indicating a low false positive rate in the barcoded assay (Fig. 3d and S8).

**Figure 4.**
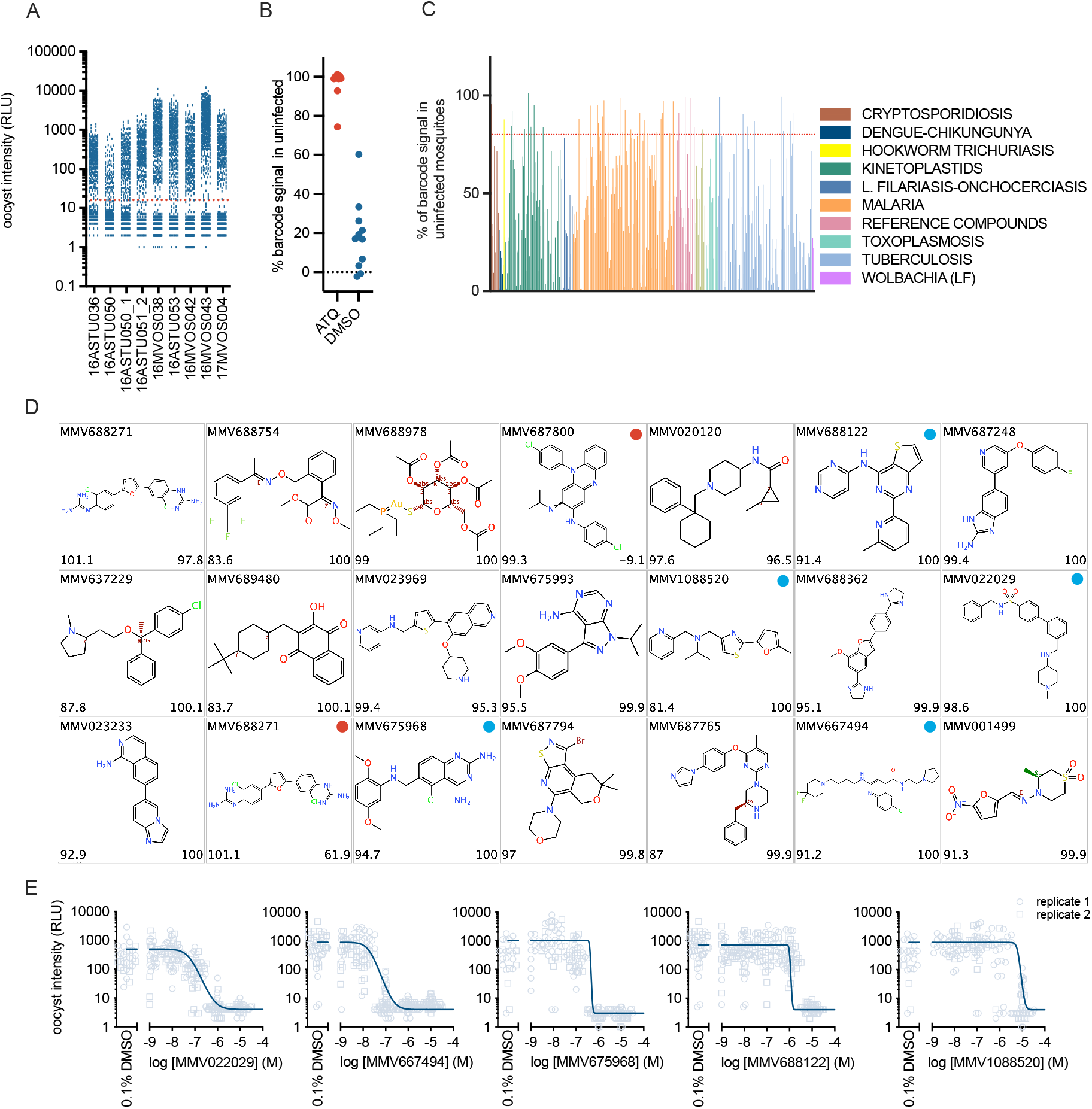
Identification of malaria transmission-blocking compounds. The open source MMV Pathogen Box was screened in a barcoded assay for *P. falciparum* transmission using a luminescent reporter parasite. Stage V gametocytes were pre-incubated with test compound at 10 or 20 μM as indicated in Table S5 and fed to *An. stephensi* mosquitoes through an *Asaia* barcoded bloodmeal. Eight days after feeding infection status was determined by a luminescence assay. Barcodes were retrieved from infected (luciferase positive) and uninfected (luciferase negative) mosquitoes and quantified. A) Oocyst intensities in mosquitoes from 9 experimental runs that were used to screen a collection of 400 compounds. The figure shows luminescence activities in individual mosquitoes. The red dotted line indicates the threshold (average background + 5σ) that was used to discriminate infected from uninfected mosquitoes. B) Assay controls. The figure shows the percentage of the total barcode signal that associated with the uninfected phenotype for barcodes in control wells containing vehicle (0.1% DMSO) or 10 μM atovaquone. C) Proportion of barcode in the uninfected mosquitoes for all compounds tested. Colors indicate the origin of the compound sets that compose the Pathogen Box. The red dotted line indicates the threshold for selection of active compounds (≥80% of the total barcode signal derived from uninfected mosquitoes). D) Compounds selected for confirmation experiments. The lower left corner of each panel indicates the proportion of barcode signal that was retrieved from uninfected mosquitoes. The lower right corner indicates the percentage reduction in oocyst intensity that was observed in a Standard Membrane Feeding Assay (SMFA). All compounds except for two compounds indicated with a red dot showed transmission blocking activity in the SMFA. Compounds identified by a blue dot were selected for full dose response analysis. E) Dose-response analysis in SMFA for selected compounds. All test concentrations were analysed in duplicate in replicate feeders. For each feeder, infection status for 24 mosquitoes was analysed through luminescence analysis. The figure shows oocyst intensities expressed as relative light units for individual mosquitoes. The solid lines indicate the fitted dose response curves.

To gain further insight in the transmission-blocking potency of a selected set of compounds, we tested 5 compounds in full dose response in glass feeder experiments. Compounds MMV1088520, MMV667494 and MMV022029 originate from the malaria compound set and the last two compounds were previously annotated as gametocytocidal (Table S6). In the dose response analyses IC_50_s were determined at 1078, 18 and 56 nM, respectively (Fig. 3e). MMV688122 originates from a *Mycobacterium* screen and blocked transmission with an IC_50_ of 1 μM whereas MMV675968 is part of the *Cryptosporidium* collection of the pathogen box and showed an IC_50_ of 406 nM. The combined results indicate that the barcoding technology significantly increases throughput in membrane feeding assays, and leads to identification of novel chemical starting points for control of malaria.

## Discussion

Conventional testing of the effectiveness of substances on longevity or vector capacity of live insects is labour intense and mostly allows only for a small number of molecules to be tested simultaneously. We have developed a technique that improves the throughput of compound testing in order to fuel pipelines for discovery of pesticides and disease transmission blocking drugs. To do this we had to overcome three distinct technical challenges: feeding mosquitoes on multiwell plates, tagging blood-fed mosquitoes with a unique well identification code, and multiplex detection of these identification codes. We used a custom designed parafilm membrane stretcher in combination with a hydraulic press to firmly seal 96 well plates filled with blood meals. The plate feeding method proved just as effective as conventional glass feeders. In order to tag mosquitoes stably throughout the course of the experiment, we used the insect midgut symbiont *Asaia* strain SF2.1, transformed with DNA-barcoded plasmids. In line with published data(*24, 34*) we observed efficient colonisation of *Anopheles* and *Aedes* mosquitoes when *Asaia* bacteria were included with the bloodmeal. Previously, Killeen *et al*. introduced a phenotypic screening concept based on phagemid encoded multisample arrays (*27*). This approach led to 95% of successfully tagged mosquitoes with the marker lasting for three days only. In contrast, *Asaia* symbionts stay with the mosquito for life and thereby makes long-term applications possible (*26*).

Our screen of a collection of pesticides exemplifies the application of the barcoding technology for discovery of novel systemic insecticides. Compounds like fluralaner and nitempyram are used as oral drugs for tick and flea control in veterinary medicine(*35, 36*) and led to enrichment of barcodes in the dead population of mosquitoes. In addition, a number of phenylpyrazoles emerged as hits with blood-borne mosquitocidal activity against *Anopheles*. Fipronil shows a very long half-life in mammalian circulation(*37*) and was shown to have potent and long-lasting mosquitocidal effects when administered to cattle(*38*). For other compounds from the phenylpyrazole class the systemic insecticide activity in a bloodmeal is less well documented, but our data show that these molecules show promise for drug-based vector control, provided they show an excellent safety profile in human. Based on the reported mammalian long *in vivo* half-life of fluralaner(*39*) this compound was selected as a promising candidate for drug-based vector control and analysed in further detail. The results, which are described elsewhere (*9*), showed potent killing activity against a wide range of vector species at concentrations that are in line with drug levels predicted to circulate for several months following a single human oral dose.

In order to exemplify a screen for vector-borne pathogen transmission, we used the barcoding technology to identify compounds that block *Plasmodium* development in *Anopheles* mosquitoes. Using the Pathogen Box collection and a selection criterion of ≥80% barcode enrichment in uninfected mosquitoes, we observed an overall hit rate of 12%. This relatively high hit rate may be explained by a biased composition of the pathogen box towards pharmacologically active compounds. A subset of 125 compounds from this collection is annotated as malaria hit compounds, as they showed IC_50_s of 2.1 μM or better against *P. falciparum* Dd2 asexual bloodstage parasites (https://www.mmv.org/mmv-open/pathogen-box). Out of these 125, 23 (18%) appear to block transmission in the barcoded screen, which is a higher number than the one predicted on basis of gametocyte viability assays(*40*). This is conceivable, as the *in vivo* transmission assay captures a wide range of potential mode of actions, including ones that incapacitate gametocytes by non-lethal ways, e.g. by prevention of gamete formation or sterilization of resulting gametes(*41*). Hit rates were 9 and 10% for compounds originating from tuberculosis and kinetoplastid hit collections that were well represented in the Pathogen Box with 116 and 70 compounds, respectively. This illustrates the strength of cross-screening bioactive molecules against a large panel of pathogen species. This notion is in line with previous observations that libraries of small molecules preselected for activity against one protozoan parasite showed high hit rates against a wider variety of pathogens(*42–44*). MMV675968 identified here as a *P. falciparum* transmission-blocking molecule belongs to a class of dihydrofolate reductase inhibitors with activity against a range of protozoa and was recently shown to block growth of *Acinetobacter baumannii*(*45, 46*). In theory, such cross-reactivity may affect the *Asaia* bacteria used in our barcoded screening strategy. As our method comprehensively monitors barcode presence in all blood-fed mosquitoes, this would lead to a total absence of the barcode in either phenotype. For the 483 compounds in the combined screens presented here we observed successful retrieval of barcode in 482 instances, indicating a relatively low hit rate against the barcode-bearing *Asaia* bacteria.

The transmission-blocking hits described here are attractive starting points for further optimisation as they obey to rule-of-five principles(*47*). In addition, all compounds have *in vitro* and *in vivo* pharmacokinetic data available (https://www.mmv.org/mmv-open/pathogen-box). For example, in rat pharmacokinetic studies, hit compound MMV687248 showed 38% absorption and clearance of 12.4 ml/min/kg, which is a reasonable starting point for further pharmacological evaluation.

Historically, phenotypic screening has driven drug R&D pipelines for infectious diseases and it has been to a larger or lesser extent been in vogue in other therapeutic areas(*48*). It is attractive as it captures complex biology in the absence of *a priori* knowledge of molecular mechanisms of disease. Recent advances in cell biological, imaging and data analyses techniques have brought it back in the spotlight(*49*). The methods described here expand the possibilities for phenotypic live insect screens. In line with published data, we observed stable colonisation of *An. stephensi* and *Ae. Aegypti* mosquitoes by *Asaia* bacteria (*24, 50*). *Asaia* has been found to associate with other sugar-feeding, phylogenetically distant genera of insects, for example the leafhopper *Scaphoideus titanus*, the vector for Flavescence Dorée, a grapevine disease (*51*). This host flexibility makes *Asaia* an attractive tool for tagging a large variety of pest insects, for the purpose of the discovery of novel pesticides and disease transmission-blocking molecules.

## Materials and methods

### Barcode Construction and transformation of *Asaia* SF2.1

Plasmid pMV170 for transformation of *Asaia* was derived by amplification of a multiple cloning site from pMV-FLPe(*52*) with primer pair MWV 371 and MWV 374 (Table S1) and introducing it into the NcoI/AatII sites of vector pBBR122 (Mobitec, Goettingen, Germany). Barcode sequences, compatible with detection using MAGPlex-TAG microspheres (Luminex Corporation, ‘s Hertogenbosch, the Netherlands) were generated by hybridization of complementary primer pairs (Table S2) and cloned into pMV170 using SpeI/AflII restriction digestion and ligation. Resulting plasmids were introduced into *E*.*coli* DH5α competent cells (ThermoFisher, Breda, the Netherlands) by heat shock transformation, yielding a collection of 50 barcoded plasmids (Table S3). Barcoded plasmids were next extracted from *E*.*coli* using the PureYield Plasmid Miniprep System (Promega, Leiden, the Netherlands) and subsequently introduced into *Asaia* sp. SF2.1 described previously(*26*). For transformation, *Asaia* cells were cultured in GLY medium (25 g/liter glycerol, 10 g/liter yeast extract, pH 5) and competent cells were prepared as previously described(*26*). Next 65 μl of the competent cells were mixed with 1 μl (∼50 ng/μl) plasmid and electroporated using a BTX electroporation system at 2.0 kV and 186 ohm in a pre-chilled 1 mm cuvette. 935 μl pre-chilled GLY medium was added and bacteria were incubated at 30°C for 4 hours without antibiotic before plating on GLY agarose plates containing 100 μg/ml kanamycin. Plates were incubated at 30 °C for 48 hours and single colonies were picked and sequence verified.

### Preparation of barcoded bloodmeals and plate feeding

Barcoded *Asaia* bacteria were grown overnight at 30°C to early log phase (OD_600_ 0.5-0.8) in a deep-well plate (Sarstedt, Nümbrecht, Germany) in 300 μl GLY medium supplemented with 100 μg/ml kanamycin per well. Bacteria were next diluted in heat inactivated human serum (type A) and combined with human red blood cells (type O) to achieve a final density of 10^6^ cfu/ml and a haematocrit of 50%. Microtiter plates were filled with 160 μl of bloodmeal per well and sealed with a membrane (Parafilm M, PM999, VWR, Amsterdam, the Netherlands) that was stretched to about 250% its original dimensions in both directions using a custom build device (Figure S1A) and applied using a lever press (Figure S1B). The plates were kept warm (37 °C) and placed upside down on top of a mosquito container sealed with mosquito netting. An aluminium block routed to fit the base of the microtiter plate and pre-heated to 45°C was put on top to warm the plate (Figure S1C). Experiments were performed with 3-5 day old females of *Anopheles stephensi* mosquitoes (Sind-Kasur Nijmegen strain) reared at the insectary of the Radboud University Medical Center(*53*), or *Aedes aegypti* (Rockefeller strain, obtained from Bayer AG, Monheim, Germany) reared at Wageningen University(*54*). For a plate containing 48 barcoded bloodmeals we used approximately 300 mosquitoes per container and for experiments with other sample sizes the number of mosquitoes was adjusted proportionally. Mosquitoes were allowed to feed for 20 minutes after which the mosquitoes were maintained at 26°C and 70-80% humidity.

### Recovery and detection of barcode sequences

Mosquitoes were washed in 70% ethanol followed by 3 washes in PBS (137 mM NaCl, 2.7 mM KCl, 10 mM Na_2_HPO_4_, 1.8 mM KH_2_PO_4_, pH 8.0). Individual mosquitoes were transferred to wells in shallow 96 well plates, combined with Zirconium beads and homogenized in 60 μl PBS using a Mini-Beadbeater-96 (Biospec, Bartlesville, OK, United States). 15 μl of each of the mosquito homogenates was subsequently transferred to a deep-well plate containing 300 μl GLY medium supplemented with 100 μg/ml kanamycin and 2 μg/ml amphotericin B. The plates were sealed with a gas permeable breathing seal (Greiner Bio-One, Alphen aan de Rijn, the Netherlands) and *Asaia* bacteria were grown to the stationary phase by incubation at 30 °C with continuous shaking (220 rpm) for at least 72 hours. In initial experiments, barcodes were amplified from individual *Asaia* cultures by PCR. In later phenotypic screening experiments, *Asaia* cultures from mosquitoes with the phenotype of interest were pooled. For comparative analyses (e.g. live versus dead mosquitoes), mock cultures with an unrelated barcode were added to make up for differences in sample sizes between the two pools. Barcodes were amplified using forward primer MWV 486 and a 5’-biotinylated reverse primer MWV 358 (Table S1) using standard PCR conditions with Gotaq G2 flexi DNA polymerase (Promega, Leiden, the Netherlands). The biotinylated PCR products were then hybridised to a pool of MagPlex-TAG microspheres (Table S3) according to manufacturer’s instructions (Luminex Corporation, ‘s Hertogenbosch, the Netherlands) with some adaptations. Briefly, 33 μl of microsphere mixture was prepared in 1.5 X TMAC hybridization solution (1x TMAC = 3M Tetramethyl ammonium chloride, 50mM Tris, 1mM EDTA and 0.1% SDS at pH 8.0) with about 1000 beads per barcode for all 50 barcode sequences. This was then mixed with 2 μl from the barcode amplification reactions and 15 μl TE buffer (10 mM Tris/1 mM EDTA, pH 8) and incubated for 15 at 52°C. Next 35 μl of reporter mix was added, containing 14.3 ug/ml SAPE (Streptavidin, R-Phycoerythrin Conjugate) and 0.24% Bovine Serum Albumin in TMAC buffer, resulting in a final concentration of 5.9 ug/ml SAPE and 0.1% BSA per reaction. After a second incubation at 52°C for 15 minutes, 50 μl was analysed on a MAGPIX instrument (Luminex Corporation, ‘s Hertogenbosch, the Netherlands).

### Screening of a collection of pesticides

A collection of pesticides was obtained through the Innovative Vector Control Consortium (Liverpool, United Kingdom) and the Medicines for Malaria Venture (Geneva, Switzerland). Compounds were first diluted in DMSO and then in human serum type A to a concentration 4 times above the final test concentration. Bloodmeals were prepared by mixing 40 μl of diluted compound with 40 μl of 4.10^6^ CFU/ml barcoded *Asaia* and 80 μl human type O red blood cells. Controls included vehicle (0.1% DMSO) and positive controls fipronil and deltamethrin, both at 10 μM. Bloodmeals were prepared in duplicate for each compound and transferred to 96 well plates in two different layouts (Figure S2). *An. stephensi* mosquitoes were allowed to feed for 20 minutes and maintained at 26°C and 70-80% humidity. 48 hours after feeding, live and dead mosquitoes were processed in separate pools as described above.

### Screening for malaria transmission-blocking compounds

Infectious *P. falciparum* gametocytes of parasite line NF54-HGL, expressing a GFP-luciferase fusion protein under control of the hsp70 promoter, were cultured in RPMI 1640 medium supplemented with 367 μM hypoxanthine, 25 mM HEPES, 25 mM sodium bicarbonate and 10% human type A serum in a semi-automated system as previously described(*20, 55*). 72 μl aliquots of cultures containing mature stage V gametocytes were transferred to 96 well v-bottom plates (Corning Life Sciences, Amsterdam, the Netherlands) in duplicate in two different layouts (Figure S2). Test compounds from the Pathogen Box (Medicines for Malaria Venture, Geneva, Switzerland) were diluted in DMSO and then in RPMI 1640 medium supplemented with 10% human serum type A and 8 μl of diluted compound was added to the gametocytes in the plate to achieve a final compound concentration of 10 or 20 μM and a final DMSO concentration of 0.2%. Positive and negative controls included 10 μM and 0.2% DMSO, respectively. Plates were incubated at 37°C, 4% CO_2_ and 3% O_2_ for 24 hours. Subsequently, plates were centrifuged briefly (750x*g*, 5’) and 70 μl supernatant was removed and replaced with 42.7 μl of heat inactivated human type A serum, 48 μl human type O red blood cells and 5.3 μl of barcoded *Asaia* bacteria to a final density of 10^5^ CFU/ml. All procedures were performed at 37 °C. Plates were then sealed and used for feeding to *An. stephensi* mosquitoes as described above. Following feeding, mosquitoes were maintained at 26°C and 70-80% humidity and starved for 2 days. From day 3 onwards, the mosquitoes were presented with cotton pads wetted in a 5% glucose solution supplemented with 100 μg/ml kanamycin twice a day for a duration of two hours each to minimize barcode cross-contamination through the glucose pads. Eight days after feeding, mosquitoes were harvested and homogenized in 96 well plates as described above. Infection status of individual mosquitoes was analyzed by determining luciferase activity in 45 μl of the mosquito homogenate as described previously(*56*). Background luminescence was determined by analyzing 10 uninfected (unfed) mosquitoes. Mosquitoes were considered infected when the luminescence signal was greater than the mean + 5xσ of the signal in the negative control mosquitoes as described previously(*20*). *Asaia* cultures were subsequently cherry picked and pooled according to infection status of the cognate mosquitoes.

### Standard Membrane Feeding Assays using glass feeders

Results from barcoded experiments were validated through Standard Membrane Feeding Assays using traditional glass feeders(*20*). For testing for systemic insecticide activity, compounds were serially diluted in DMSO and then in DMEM medium and combined with human type A serum and type O red blood cells to achieve a final DMSO concentration of 0.1% in 40% haematocrit in a volume of 300 μl. Bloodmeals were placed in glass feeders warmed at 37°C and *An. stephensi* mosquitoes were allowed to feed for 15 minutes. Following feeding, non-fed mosquitoes were removed and the blood-fed mosquitoes were maintained at 26°C and 70-80% humidity for 48 hours. Subsequently, the number of live and dead mosquitoes was determined for each test condition. Testing for compound effects on transmission of *P. falciparum* gametocytes to *An. stephensi* mosquitoes was performed as described previously(*20*).

### Replicates and data analyses

To obtain sufficient numbers of fed mosquitoes, all test compounds were presented in replicate bloodmeals (Figure S2). An average of 6 mosquitoes per bloodmeal was used in barcoded feeding experiments. With a 90% feeding efficiency, this resulted in ∼10 fed mosquitoes per test condition. Mosquitoes were processed individually and rescued barcoded *Asaia* bacteria were pooled according to phenotype. Here, the *Asaia* from the replicate plates were combined for each phenotype. For each pool, barcode fragments were amplified and analysed in triplicate. Fluorescence intensity was determined by analyses of at least 40 microspheres per barcode and expressed as relative median fluorescence intensity (MFI). MFI values were averaged from the triplicates observations for each pool and corrected for average background signals from negative control (GLY-medium without barcoded *Asaia*) samples. Barcodes were considered as sampled when the signal was above the mean + 3σ of the negative control samples. In comparative phenotypic analyses, data were expressed as the relative proportion of the barcode signal in the phenotype of interest. In Standard Membrane Feeding Experiments using glass feeders, all conditions were tested in two replicate feeders, and at least 24 mosquitoes were analysed per feeder. Data were analysed and visualised using the GraphPad Prism software package. IC_50_ values for systemic insecticides were determined by fitting a four parameter logistic regression model using least squares to find the best fit. IC_50_ values in *Plasmodium* transmission blocking experiments were determined by assuming a beta binomial distribution and logistic regression using maximum likelihood to find the best fit as described previously(*57*). Effects of Pantoea or Asaia on *P. falciparum* were analysed by ANOVA using a Kruskall-Wallis test and Dunn’s multiple comparison test.

## Acknowledgements

We wish to thank Claudia Damiani and Aida Capone for help with the *Asaia* SF2.1 strain, Marcelo Jacobs-Lorena for his kind gift of *Pantoea agglomerans* and Sander Koenraadt for provision of *Aedes aegypti* mosquitoes. Bernd Engelbrecht, Katharina Schumacher and colleagues at Irmato Industrial Solutions are gratefully acknowledged for help with the design of the plate sealing process. The authors wish to thank Geert-Jan van Gemert and Laura Pelsen-Posthumus for expert technical assistance in mosquito rearing. Sarah Rees is acknowledged for provision of a collection of pesticides. The authors thank Isaac Sandoval Capuchino for providing artwork. Manuel Llinás and Robert Sauerwein are gratefully acknowledged for critical reading of the manuscript. This work was supported, in whole or in part, by the Bill & Melinda Gates Foundation (grants OPP1067662 & OPP1118462). Under the grant conditions of the Foundation, a Creative Commons Attribution 4.0 Generic License has already been assigned to the Author Accepted Manuscript version that might arise from this submission.

## Author contributions

GF, KS, EH and KJD conceived the work and supervised experiments. AS, MWV, RH, ME, KMJK, AvS generated and analyzed data. GF contributed novel technologies and reagents. AS, MWV and KJD drafted the manuscript. All authors proof-read and edited the manuscript.

Conceptualization and supervision: KS, EH and KJD

Novel technologies and reagents: GF

Experimentation and data analyses: ASt, MWV, RH, ME, KMJK, ASh

Writing – original draft: ASt and MWV

Writing – review & editing: ASt and KJD

## References

1. S. Bhatt, D. J. Weiss, E. Cameron, D. Bisanzio, B. Mappin, U. Dalrymple, K. E. Battle, C. L. Moyes, a. Henry, P. a. Eckhoff, E. a. Wenger, O. Briët, M. a. Penny, T. a. Smith, a. Bennett, J. Yukich, T. P. Eisele, J. T. Griffin, C. a. Fergus, M. Lynch, F. Lindgren, J. M. Cohen, C. L. J. Murray, D. L. Smith, S. I. Hay, R. E. Cibulskis, P. W. Gething, The effect of malaria control on Plasmodium falciparum in Africa between 2000 and 2015. Nature (2015), doi:10.1038/nature15535.

2. D. R. Swale, Perspectives on new strategies for the identification and development of insecticide targets. Pesticide Biochemistry and Physiology. 161, 23–32 (2019).

3. T. C. Sparks, R. Nauen, IRAC : Mode of action classification and insecticide resistance management. Pesticide Biochemistry and Physiology. 121, 122–128 (2015).

4. J. Burrows, H. Slater, F. Macintyre, S. Rees, A. Thomas, F. Okumu, R. Hooft Van Huijsduijnen, S. Duparc, T. N. C. Wells, A discovery and development roadmap for new endectocidal transmission-blocking agents in malaria. Malaria Journal. 17, 1–15 (2018).

5. C. Chaccour, J. Lines, C. J. M. Whitty, Effect of ivermectin on Anopheles gambiae mosquitoes fed on humans: the potential of oral insecticides in malaria control. The Journal of infectious diseases. 202, 113–6 (2010).

6. B. D. Foy, H. Alout, J. A. Seaman, S. Rao, T. Magalhaes, M. Wade, S. Parikh, D. D. Soma, A. B. Sagna, F. Fournet, H. C. Slater, R. Bougma, F. Drabo, A. Diabaté, A. G. v. Coulidiaty, N. Rouamba, R. K. Dabiré, Efficacy and risk of harms of repeat ivermectin mass drug administrations for control of malaria (RIMDAMAL): a cluster-randomised trial. The Lancet. 393, 1517–1526 (2019).

7. H. C. Slater, P. G. T. Walker, T. Bousema, L. C. Okell, A. C. Ghani, The potential impact of adding ivermectin to a mass treatment intervention to reduce malaria transmission: a modelling study. The Journal of infectious diseases. 210, 1972–80 (2014).

8. A. M. Bellinger, M. Jafari, T. M. Grant, S. Zhang, H. C. Slater, E. A. Wenger, S. Mo, Y.-A. L. Lee, H. Mazdiyasni, L. Kogan, R. Barman, C. Cleveland, L. Booth, T. Bensel, D. Minahan, H. M. Hurowitz, T. Tai, J. Daily, B. Nikolic, L. Wood, P. A. Eckhoff, R. Langer, G. Traverso, Oral, ultra-long-lasting drug delivery: Application toward malaria elimination goals. Science translational medicine. 8, 365ra157 (2016).

9. M. Miglianico, M. Eldering, H. Slater, N. Ferguson, P. Ambrose, R. S. Lees, K. M. J. J. Koolen, K. Pruzinova, M. Jancarova, P. Volf, C. J. M. M. Koenraadt, H.-P. P. Duerr, G. Trevitt, B. Yang, A. K. Chatterjee, J. Wisler, A. Sturm, T. Bousema, R. W. Sauerwein, P. G. Schultz, M. S. Tremblay, K. J. Dechering, Repurposing isoxazoline veterinary drugs for control of vector-borne human diseases. Proceedings of the National Academy of Sciences of the United States of America. 115, E6920–E6926 (2018).

10. M. A. Phillips, J. N. Burrows, C. Manyando, R. H. van Huijsduijnen, W. C. van Voorhis, T. N. C. Wells, Malaria (2017), doi:10.1038/nrdp.2017.50.

11. P. M. Armstrong, H. Y. Ehrlich, T. Magalhaes, M. R. Miller, P. J. Conway, A. Bransfield, M. J. Misencik, A. Gloria-Soria, J. L. Warren, T. G. Andreadis, J. J. Shepard, B. D. Foy, V. E. Pitzer, D. E. Brackney, Successive blood meals enhance virus dissemination within mosquitoes and increase transmission potential. Nature Microbiology (2019), doi:10.1038/s41564-019-0619-y.

12. S. Yahiya, A. Rueda-Zubiaurre, M. J. Delves, M. J. Fuchter, J. Baum, The antimalarial screening landscape—looking beyond the asexual blood stage. Current Opinion in Chemical Biology. 50, 1–9 (2019).

13. D. M. Plouffe, M. Wree, A. Y. Du, C. A. Scherer, J. Vinetz, E. A. Winzeler, D. M. Plouffe, M. Wree, A. Y. Du, S. Meister, F. Li, K. Patra, A. Lubar, High-Throughput Assay and Discovery of Small Molecules that Interrupt Malaria Transmission Resource High-Throughput Assay and Discovery of Small Molecules that Interrupt Malaria Transmission. Cell Host and Microbe, 1–13 (2016).

14. C. Miguel-Blanco, I. Molina, A. I. Bardera, B. Díaz, L. de las Heras, S. Lozano, C. González, J. Rodrigues, M. J. Delves, A. Ruecker, G. Colmenarejo, S. Viera, M. S. Martínez-Martínez, E. Fernández, J. Baum, R. E. Sinden, E. Herreros, Hundreds of dual-stage antimalarial molecules discovered by a functional gametocyte screen. Nature Communications. 8, 15160 (2017).

15. K. J. K. J. K. J. Dechering, H.-P. H.-P. P. Duerr, K. M. J. K. M. J. J. Koolen, G.-J. J. V. G.-J. V. Gemert, T. Bousema, J. Burrows, D. Leroy, R. W. R. W. Sauerwein, Modelling mosquito infection at natural parasite densities identifies drugs targeting EF2, PI4K or ATP4 as key candidates for interrupting malaria transmission. Scientific Reports. 7, 17680 (2017).

16. J. Schalkwijk, E. L. E. L. Allman, P. A. M. M. P. A. M. Jansen, L. E. L. E. de Vries, J. M. J. J. M. J. J. Verhoef, S. Jackowski, P. N. M. P. N. M. M. Botman, C. A. C. A. Beuckens-Schortinghuis, K. M. J. K. M. J. J. Koolen, J. M. J. M. J. M. Bolscher, M. W. M. W. Vos, K. Miller, S. A. Reeves, H. Pett, G. Trevitt, S. Wittlin, C. Scheurer, S. Sax, C. Fischli, I. I. Angulo-Barturen, M. B. M. B. Jiménez-Diaz, G. Josling, T. W. A. A. T. W. A. Kooij, R. Bonnert, B. Campo, R. H. R. H. Blaauw, F. P. J. T. F. P. J. T. F. P. J. T. Rutjes, R. W. R. W. Sauerwein, M. Llinás, P. H. H. P. H. H. H. Hermkens, K. J. K. J. Dechering, M. B. Jimenez-Diaz, G. Josling, T. W. A. A. T. W. A. Kooij, R. Bonnert, B. Campo, R. H. R. H. Blaauw, F. P. J. T. F. P. J. T. F. P. J. T. Rutjes, R. W. R. W. Sauerwein, M. Llinas, P. H. H. P. H. H. H. Hermkens, K. J. K. J. Dechering, Antimalarial pantothenamide metabolites target acetyl-coenzyme A biosynthesis in Plasmodium falciparum. Science Translational Medicine. 11 (2019), doi:10.1126/scitranslmed.aas9917.

17. J. N. Burrows, S. Duparc, W. E. Gutteridge, R. Hooft van Huijsduijnen, W. Kaszubska, F. Macintyre, S. Mazzuri, J. J. Möhrle, T. N. C. Wells, New developments in anti-malarial target candidate and product profiles. Malaria Journal. 16, 26 (2017).

18. T. Yang, S. Ottilie, E. S. Istvan, K. P. Godinez-Macias, A. K. Lukens, B. Baragaña, B. Campo, C. Walpole, J. C. Niles, K. Chibale, K. J. Dechering, M. Llinás, M. C. S. Lee, N. Kato, S. Wyllie, C. W. McNamara, F. J. Gamo, J. Burrows, D. A. Fidock, D. E. Goldberg, H. Gilbert, D. F. Wirth, E. A. Winzeler, A. Malaria Drug Accelerator Consortium, MalDA, Accelerating Malaria Drug Discovery. Trends in parasitology. 37, 493–507 (2021).

19. J. A. Turner, C. N. E. Ruscoe, T. R. Perrior, Discovery to Development : Insecticides for Malaria Vector Control. 70, 684–693 (2016).

20. M. W. Vos, W. J. R. Stone, K. M. Koolen, G.-J. van Gemert, B. van Schaijk, D. Leroy, R. W. Sauerwein, T. Bousema, K. J. Dechering, A semi-automated luminescence based standard membrane feeding assay identifies novel small molecules that inhibit transmission of malaria parasites by mosquitoes. Scientific Reports. 5 (2015), doi:10.1038/srep18704.

21. T. C. Sparks, F. J. Wessels, B. A. Lorsbach, B. M. Nugent, G. B. Watson, The new age of insecticide discovery-the crop protection industry and the impact of natural products. Pesticide Biochemistry and Physiology. 161, 12–22 (2019).

22. S. D. Buckingham, F. A. Partridge, B. C. Poulton, B. S. Miller, R. A. McKendry, G. J. Lycett, D. B. Sattelle, Automated phenotyping of mosquito larvae enables high-throughput screening for novel larvicides and offers potential for smartphone-based detection of larval insecticide resistance. PLoS neglected tropical diseases. 15, e0008639 (2021).

23. K. J. Dechering, H.-P. Duerr, K. M. J. Koolen, G.-J. V. Gemert, T. Bousema, J. Burrows, D. Leroy, R. W. Sauerwein, Modelling mosquito infection at natural parasite densities identifies drugs targeting EF2, PI4K or ATP4 as key candidates for interrupting malaria transmission. Scientific Reports. 7 (2017), doi:10.1038/s41598-017-16671-0.

24. E. Crotti, C. Damiani, M. Pajoro, E. Gonella, A. Rizzi, I. Ricci, I. Negri, P. Scuppa, P. Rossi, P. Ballarini, N. Raddadi, M. Marzorati, L. Sacchi, E. Clementi, M. Genchi, M. Mandrioli, C. Bandi, G. Favia, A. Alma, D. Daffonchio, Asaia, a versatile acetic acid bacterial symbiont, capable of cross-colonizing insects of phylogenetically distant genera and orders. Environmental Microbiology. 11, 3252–3264 (2009).

25. F. Li, P. Li, H. Hua, M. Hou, F. Wang, Diversity, Tissue Localization, and Infection Pattern of Bacterial Symbionts of the White-Backed Planthopper, Sogatella furcifera (Hemiptera: Delphacidae). Microbial Ecology (2019), doi:10.1007/s00248-019-01433-4.

26. G. Favia, I. Ricci, C. Damiani, N. Raddadi, E. Crotti, M. Marzorati, A. Rizzi, R. Urso, L. Brusetti, S. Borin, D. Mora, P. Scuppa, L. Pasqualini, E. Clementi, M. Genchi, S. Corona, M. Negri, G. Grandi, A. Alma, L. Kramer, F. Esposito, C. Bandi, L. Sacchi, D. Daffonchio, Bacteria of the genus Asaia stably associate with Anopheles stephensi, an Asian malarial mosquito vector. Proceedings of the National Academy of Sciences of the United States of America. 104, 9047–9051 (2007).

27. G. F. Killeen, B. D. Foy, M. Shahabuddin, W. Roake, a Williams, T. J. Vaughan, J. C. Beier, Tagging bloodmeals with phagemids allows feeding of multiple-sample arrays to single cages of mosquitoes (Diptera: Culicidae) and the recovery of single recombinant antibody fragment genes from individual insects. Journal of medical entomology. 37, 528–533 (2000).

28. S. A. Dunbar, Applications of Luminex^®^ xMAP™ technology for rapid, high-throughput multiplexed nucleic acid detection. Clinica Chimica Acta. 363, 71–82 (2006).

29. S. Wang, a. K. Ghosh, N. Bongio, K. a. Stebbings, D. J. Lampe, M. Jacobs-Lorena, Fighting malaria with engineered symbiotic bacteria from vector mosquitoes. Proceedings of the National Academy of Sciences. 109, 12734–12739 (2012).

30. M. W. M. W. Vos, W. J. R. W. J. R. Stone, K. M. K. M. K. M. Koolen, G.-J. G.-J. van Gemert, B. van Schaijk, D. Leroy, R. W. R. W. Sauerwein, T. Bousema, K. J. K. J. K. J. Dechering, A semi-automated luminescence based standard membrane feeding assay identifies novel small molecules that inhibit transmission of malaria parasites by mosquitoes. Scientific Reports. 5, 18704 (2015).

31. W. J. R. W. J. R. J. Stone, T. S. T. S. Churcher, W. Graumans, G.-J. G.-J. J. van Gemert, M. W. M. W. Vos, K. H. W. K. H. W. Lanke, M. G. M. G. G. van de Vegte-Bolmer, R. Siebelink-Stoter, K. J. K. J. J. Dechering, A. M. M. A. M. Vaughan, N. Camargo, S. H. I. S. H. I. Kappe, R. W. R. W. Sauerwein, T. Bousema, A Scalable Assessment of Plasmodium falciparum Transmission in the Standard Membrane-Feeding Assay, Using Transgenic Parasites Expressing Green Fluorescent Protein-Luciferase. Journal of Infectious Diseases. 210, 1456–1463 (2014).

32. C. G. L. Veale, Unpacking the Pathogen Box—An Open Source Tool for Fighting Neglected Tropical Disease. ChemMedChem. 14, 386–453 (2019).

33. T. S. Churcher, A. M. Blagborough, M. Delves, C. Ramakrishnan, M. C. Kapulu, A. R. Williams, S. Biswas, D. F. Da, A. Cohuet, R. E. Sinden, Measuring the blockade of malaria transmission--an analysis of the Standard Membrane Feeding Assay. International Journal for Parasitology. 42, 1037–1044 (2012).

34. C. Damiani, I. Ricci, E. Crotti, P. Rossi, A. Rizzi, P. Scuppa, A. Capone, U. Ulissi, S. Epis, M. Genchi, N. Sagnon, I. Faye, A. Kang, B. Chouaia, C. Whitehorn, G. W. Moussa, M. Mandrioli, F. Esposito, L. Sacchi, C. Bandi, D. Daffonchio, G. Favia, Mosquito-bacteria symbiosis: the case of Anopheles gambiae and Asaia. Microbial Ecology. 60, 644–654 (2010).

35. C. Wengenmayer, H. Williams, E. Zschiesche, A. Moritz, J. Langenstein, R. K. A. Roepke, A. R. Heckeroth, The speed of kill of fluralaner (Bravecto ™) against Ixodes ricinus ticks on dogs, 1–5 (2014).

36. R. Schenker, O. Tinembart, E. Humbert-Droz, T. Cavaliero, B. Yerly, Comparative speed of kill between nitenpyram, fipronil, imidacloprid, selamectin and cythioate against adult Ctenocephalides felis (Bouché) on cats and dogs. Veterinary Parasitology. 112, 249–254 (2003).

37. G. C. M. dos Santos, L. H. G. Rosado, M. C. C. Alves, I. de Paula Lima, T. P. Ferreira, D. A. Borges, P. C. de Oliveira, V. de Sousa Magalhães, F. B. Scott, Y. P. Cid, Fipronil Tablets: Development and Pharmacokinetic Profile in Beagle Dogs. AAPS PharmSciTech. 21, 9 (2020).

38. R. M. Poché, N. Githaka, F. van Gool, R. C. Kading, D. Hartman, L. Polyakova, E. O. Abworo, V. Nene, S. Lozano-Fuentes, Preliminary efficacy investigations of oral fipronil against Anopheles arabiensis when administered to Zebu cattle (Bos indicus) under field conditions. Acta Tropica. 176, 126–133 (2017).

39. S. Kilp, D. Ramirez, M. J. Allan, R. K. a Roepke, M. C. Nuernberger, Pharmacokinetics of fluralaner in dogs following a single oral or intravenous administration. Parasites & vectors. 7, 85 (2014).

40. S. Duffy, M. L. Sykes, A. J. Jones, T. B. Shelper, M. Simpson, R. Lang, S.-A. Poulsen, B. E. Sleebs, V. M. Avery, Screening the Medicines for Malaria Venture Pathogen Box across Multiple Pathogens Reclassifies Starting Points for Open-Source Drug Discovery. Antimicrobial agents and chemotherapy. 61 (2017), doi:10.1128/AAC.00379-17.

41. a. Ruecker, D. K. Mathias, U. Straschil, T. S. Churcher, R. R. Dinglasan, D. Leroy, R. E. Sinden, M. J. Delves, A Male and Female Gametocyte Functional Viability Assay To Identify Biologically Relevant Malaria Transmission-Blocking Drugs. Antimicrobial Agents and Chemotherapy. 58, 7292–7302 (2014).

42. W. C. van Voorhis, J. H. Adams, R. Adelfio, V. Ahyong, M. H. Akabas, P. Alano, A. Alday, Y. Alemán Resto, A. Alsibaee, A. Alzualde, K. T. Andrews, S. v. Avery, V. M. Avery, L. Ayong, M. Baker, S. Baker, C. ben Mamoun, S. Bhatia, Q. Bickle, L. Bounaadja, T. Bowling, J. Bosch, L. E. Boucher, F. F. Boyom, J. Brea, M. Brennan, A. Burton, C. R. Caffrey, G. Camarda, M. Carrasquilla, D. Carter, M. Belen Cassera, K. Chih-Chien Cheng, W. Chindaudomsate, A. Chubb, B. L. Colon, D. D. Colón-López, Y. Corbett, G. J. Crowther, N. Cowan, S. D’Alessandro, N. le Dang, M. Delves, J. L. DeRisi, A. Y. Du, S. Duffy, S. Abd El-Salam El-Sayed, M. T. Ferdig, J. A. Fernández Robledo, D. A. Fidock, I. Florent, P. V. T. Fokou, A. Galstian, F. J. Gamo, S. Gokool, B. Gold, T. Golub, G. M. Goldgof, R. Guha, W. A. Guiguemde, N. Gural, R. K. Guy, M. A. E. Hansen, K. K. Hanson, A. Hemphill, R. Hooft van Huijsduijnen, T. Horii, P. Horrocks, T. B. Hughes, C. Huston, I. Igarashi, K. Ingram-Sieber, M. A. Itoe, A. Jadhav, A. Naranuntarat Jensen, L. T. Jensen, R. H. Y. Jiang, A. Kaiser, J. Keiser, T. Ketas, S. Kicka, S. Kim, K. Kirk, V. P. Kumar, D. E. Kyle, M. J. Lafuente, S. Landfear, N. Lee, S. Lee, A. M. Lehane, F. Li, D. Little, L. Liu, M. Llinás, M. I. Loza, A. Lubar, L. Lucantoni, I. Lucet, L. Maes, D. Mancama, N. R. Mansour, S. March, S. McGowan, I. Medina Vera, S. Meister, L. Mercer, J. Mestres, A. N. Mfopa, R. N. Misra, S. Moon, J. P. Moore, F. Morais Rodrigues da Costa, J. Müller, A. Muriana, S. Nakazawa Hewitt, B. Nare, C. Nathan, N. Narraidoo, S. Nawaratna, K. K. Ojo, D. Ortiz, G. Panic, G. Papadatos, S. Parapini, K. Patra, N. Pham, S. Prats, D. M. Plouffe, S. A. Poulsen, A. Pradhan, C. Quevedo, R. J. Quinn, C. A. Rice, M. Abdo Rizk, A. Ruecker, R. st. Onge, R. Salgado Ferreira, J. Samra, N. G. Robinett, U. Schlecht, M. Schmitt, F. Silva Villela, F. Silvestrini, R. Sinden, D. A. Smith, T. Soldati, A. Spitzmüller, S. M. Stamm, D. J. Sullivan, W. Sullivan, S. Suresh, B. M. Suzuki, Y. Suzuki, S. J. Swamidass, D. Taramelli, L. R. Y. Tchokouaha, A. Theron, D. Thomas, K. F. Tonissen, S. Townson, A. K. Tripathi, V. Trofimov, K. O. Udenze, I. Ullah, C. Vallieres, E. Vigil, J. M. Vinetz, P. Voong Vinh, H. Vu, N. A. Watanabe, K. Weatherby, P. M. White, A. F. Wilks, E. A. Winzeler, E. Wojcik, M. Wree, W. Wu, N. Yokoyama, P. H. A. Zollo, N. Abla, B. Blasco, J. Burrows, B. Laleu, D. Leroy, T. Spangenberg, T. Wells, P. A. Willis, Open Source Drug Discovery with the Malaria Box Compound Collection for Neglected Diseases and Beyond. PLoS Pathogens. 12, 1–23 (2016).

43. W. Devine, J. L. Woodring, U. Swaminathan, E. Amata, G. Patel, J. Erath, N. E. Roncal, P. J. Lee, S. E. Leed, A. Rodriguez, K. Mensa-Wilmot, R. J. Sciotti, M. P. Pollastri, Protozoan Parasite Growth Inhibitors Discovered by Cross-Screening Yield Potent Scaffolds for Lead Discovery. Journal of Medicinal Chemistry. 58, 5522–5537 (2015).

44. J. L. Woodring, K. A. Bachovchin, K. G. Brady, M. F. Gallerstein, J. Erath, S. Tanghe, S. E. Leed, A. Rodriguez, K. Mensa-Wilmot, R. J. Sciotti, M. P. Pollastri, Optimization of physicochemical properties for 4-anilinoquinazoline inhibitors of trypanosome proliferation. European Journal of Medicinal Chemistry. 141, 446–459 (2017).

45. W. Songsungthong, S. Yongkiettrakul, L. E. Bohan, E. S. Nicholson, S. Prasopporn, P. Chaiyen, U. Leartsakulpanich, Diaminoquinazoline MMV675968 from Pathogen Box inhibits Acinetobacter baumannii growth through targeting of dihydrofolate reductase. Scientific Reports. 9, 15625 (2019).

46. H. Lau, J. T. Ferlan, V. H. Brophy, A. Rosowsky, C. H. Sibley, Efficacies of lipophilic inhibitors of dihydrofolate reductase against parasitic protozoa. Antimicrobial agents and chemotherapy. 45, 187–95 (2001).

47. C. A. Lipinski, F. Lombardo, B. W. Dominy, P. J. Feeney, Experimental and computational approaches to estimate solubility and permeability in drug discovery and development settings. Advanced Drug Delivery Reviews. 46, 3–26 (2001).

48. W. Zheng, N. Thorne, J. C. McKew, Phenotypic screens as a renewed approach for drug discovery. Drug discovery today. 18, 1067–73 (2013).

49. N. Aulner, A. Danckaert, J. Ihm, D. Shum, S. L. Shorte, Next-Generation Phenotypic Screening in Early Drug Discovery for Infectious Diseases. Trends in Parasitology. 35, 559–570 (2019).

50. A. Capone, I. Ricci, C. Damiani, M. Mosca, P. Rossi, P. Scuppa, E. Crotti, S. Epis, M. Angeletti, M. Valzano, L. Sacchi, C. Bandi, D. Daffonchio, M. Mandrioli, G. Favia, Interactions between Asaia, Plasmodium and Anopheles: new insights into mosquito symbiosis and implications in malaria symbiotic control. Parasites & vectors. 6, 182 (2013).

51. E. Crotti, M. Pajoro, C. Damiani, I. Ricci, I. Negri, A. Rizzi, E. Clementi, N. Raddadi, P. Scuppa, M. Marzorati, L. Pasqualini, C. Bandi, L. Sacchi, G. Favia, A. Alma, D. Daffonchio, Asaia, a transformable bacterium, associated with Scaphoideus titanus, the vector of “flavescence dorée.” Bulletin of Insectology. 61, 219–220 (2008).

52. B. C. van Schaijk, M. W. Vos, C. J. Janse, R. W. Sauerwein, S. M. Khan, Removal of heterologous sequences from Plasmodium falciparum mutants using FLPe-recombinase. PloS One. 5, e15121 (2010).

53. A. M. Feldmann, T. Ponnudurai, Selection of Anopheles stephensi for refractoriness and susceptibility to Plasmodium falciparum. Medical and Veterinary Entomology. 3, 41–52 (1989).

54. G. P. Göertz, C. B. F. Vogels, C. Geertsema, C. J. M. Koenraadt, G. P. Pijlman, Mosquito co-infection with Zika and chikungunya virus allows simultaneous transmission without affecting vector competence of Aedes aegypti. PLOS Neglected Tropical Diseases. 11, e0005654 (2017).

55. T. Ponnudurai, A. H. Lensen, G. J. van Gemert, M. P. Bensink, M. Bolmer, J. H. Meuwissen, Infectivity of cultured Plasmodium falciparum gametocytes to mosquitoes. Parasitology. 98 Pt 2, 165–73 (1989).

56. W. J. R. Stone, T. S. Churcher, W. Graumans, G.-J. van Gemert, M. W. Vos, K. H. W. Lanke, M. G. van de Vegte-Bolmer, R. Siebelink-Stoter, K. J. Dechering, A. M. Vaughan, N. Camargo, S. H. I. Kappe, R. W. Sauerwein, T. Bousema, A scalable assessment of Plasmodium falciparum transmission in the standard membrane-feeding assay, using transgenic parasites expressing green fluorescent protein-luciferase. Journal of Infectious Diseases. 210 (2014), doi:10.1093/infdis/jiu271.

57. K. J. Dechering, H. P. Duerr, K. M. J. Koolen, G. J. van Gemert, T. Bousema, J. Burrows, D. Leroy, R. W. Sauerwein, Modelling mosquito infection at natural parasite densities identifies drugs targeting EF2, PI4K or ATP4 as key candidates for interrupting malaria transmission. Scientific Reports. 7, 1–9 (2017).

